# A Novel Dual Probe-based Method for Mutation Detection using Isothermal Amplification

**DOI:** 10.1101/2024.03.27.586979

**Authors:** Nidhi Nandu, Michael Miller, Yanhong Tong, Zhi-xiang Lu

## Abstract

Cost efficient and rapid detection tools to detect mutations especially those linked to drug-resistance are important to address concerns of the rising multi-drug resistance infections. Here we integrated dual probes, namely a calibrator probe and an indicator probe, into isothermal amplification detection system. These two probes are designed to bind distinct regions on the same amplicon to determine the presence or absence of mutation. The calibrator probe signal is used as an internal signal calibrator for indicator probe which detects the presence or absence of the mutation. As an illustrative example, we evaluated the applicability of this dual probe method for detecting mutations associated with rifampicin (RIF) drug resistance at codons 516, 526 and 531 of the rpoB gene in *Mycobacterium tuberculosis*. In this assessment, we examined 127 artificial samples comprising wild type and mutant target sequences with single or multiple mutations. Our results demonstrated 100% accuracy of both wild type and mutant samples for mutations at codons 526 and 531. As regards to mutations at codon 516, the wild type was identified with 100% accuracy, while the mutant type was identified with 95% accuracy. Moreover, when we extended our evaluation using the Zeptometrix MTB Verification panel, our dual probe method correctly differentiated between the wild type and mutant, and identified the RIF-mutant strain which harbors mutations at codon 531 of the rpoB gene. Our isothermal mutation detection system, relying on dual probes exhibits a versatile approach. With the capability to identify mutations without prior knowledge of their specific mutation direction, our dual-probe method shows significant promise for applications in drug resistance nucleic acid testing, particularly in resource-limited settings.

## INTRODUCTION

Mutations are broadly a result of random glitches in DNA replicating machinery which play an important role in evolution. Unfortunately, the golden era of antibiotics and non-judicious use of antibiotics, contributed to selective rise of mutations conferring drug-resistance. Modern medicine which relies on using antibiotics for curing infections is currently threatened by the emergence of antibiotic resistant strains. Traditionally, drug susceptibility testing (DST) methods, such as phenotypic culture-based assays and more recently molecular techniques have been employed for resistance detection. Polymerase chain reaction (PCR) based methods, such as ARMS-PCR, CLAMP-PCR, or hydrolysis probe Melting Curve Analysis, primer extension SNaPshot analysis after PCR amplification, are widely used to detect mutations in research and diagnostic applications.^1,2^ However, the need for thermocyclers limits the application of these methods. In general, many of these methods are often time-consuming, require sophisticated laboratory facilities and trained personnel, and may be expensive, making them less suitable for resource-limited settings where the burden of infectious disease is the highest. This has led to the development of many alternative technologies like isothermal nucleic acid amplification technologies for mutation detection that can be carried out at a constant temperature eliminating the need for sophisticated thermal-cycling equipment. Especially, in cases where the aim is not to quantify infection, but to confirm the presence or absence of infection and drug-resistance, the rapid turnaround time can be helpful in guiding patient treatment plans and decrease instances of unnecessary use of antibiotics. These methods can aid in early detection of drug-resistance infection spread and contribute towards fight against growing drug-resistance.

Many of the vastly researched isothermal techniques suffer from certain disadvantages that limit their application for mutation detection. The properties of enzymes or polymerases used in isothermal reaction along with no temperature cycling for renaturing and denaturing of DNA, lead to reduced stringency of primer annealing and extension which result in isothermal technologies tolerating mismatches or mutations in test target.^3,4^ Although researchers have demonstrated the feasibility of using isothermal amplification technology by integrating endonuclease (RNase H2 or T^th^ endonuclease IV), via ligation-initiation, or using self-stabilizing competitive primers for mutation detection, these novel methods require complicated ingredients. Moreover, some of these methods are limited to detect only one mutation per reaction, and some of the methods require multiple primer sets to detect a single mutation.^5–9^

Tuberculosis (TB) is one such disease, that still haunts us in the present due to the emerging multi-drug resistant (MDR) strains. As a major global health threat TB causes millions of new infections and deaths each year. As recent as 2020, compounded by the COVID-19 pandemic, an estimated 1.9 million deaths were attributed to tuberculosis.^10^ The global efforts to fight TB are continuously being thwarted by the emerging drug-resistance strains, particularly those resistant to rifampicin (RIF) pose a significant challenge.^11^ Rifampicin is a key first-line antibiotic used in the standard TB treatment regimen, and resistance to this drug combined with isoniazid resistance is often a marker of multidrug-resistant TB (MDR-TB) or extensively drug-resistant TB (XDR-TB).^12^ Prompt and accurate detection of RIF resistance (RR-TB) is crucial for guiding appropriate treatment decisions, preventing the spread of drug-resistant strains, and improving patient outcomes. Mutations commonly occurring in the 81-bp long region of the *rpoB* gene, which encodes the beta subunit of RNA polymerase, called the RIF resistance-determining region (RRDR) can lead to resistance to RIF (also referred to as RIF-resistance). Ninety percent of the RIF-resistance is linked to mutations in this key region, with most common mutations occurring at codons 516 (resulting in Asp to Val or Tyr), 526 (resulting in His to Leu or Tyr), and 531(resulting in Ser to Leu or Trp) based on *E. coli rpoB* gene numbering system.^13,14^

In this study, we established a dual probe method for mutation detection. To demonstrate its applicability in resource limited setting, we selected Loop mediated amplification (LAMP) to amplify the target sequence. This dual probe method design allows for use of two probes that bind to different regions of the same amplicon formed from a primer set making the assay design simple. One of the probes binds to a conserved sequence acting as a signal calibrator (Probe C) and the second probe binds to the mutation site acting as a mutation indicator (Probe I) (Figure 1). Additionally, considering signals from both the probes helps mitigate variations introduced due to reaction conditions. We also show that multiple closely located mutations can be detected using same primer set. Although the dual probe method described in this study was used in conjunction with LAMP isothermal amplification, the dual probe strategy can be applied to amplicons generated by any other isothermal technologies.

**Figure 1:**
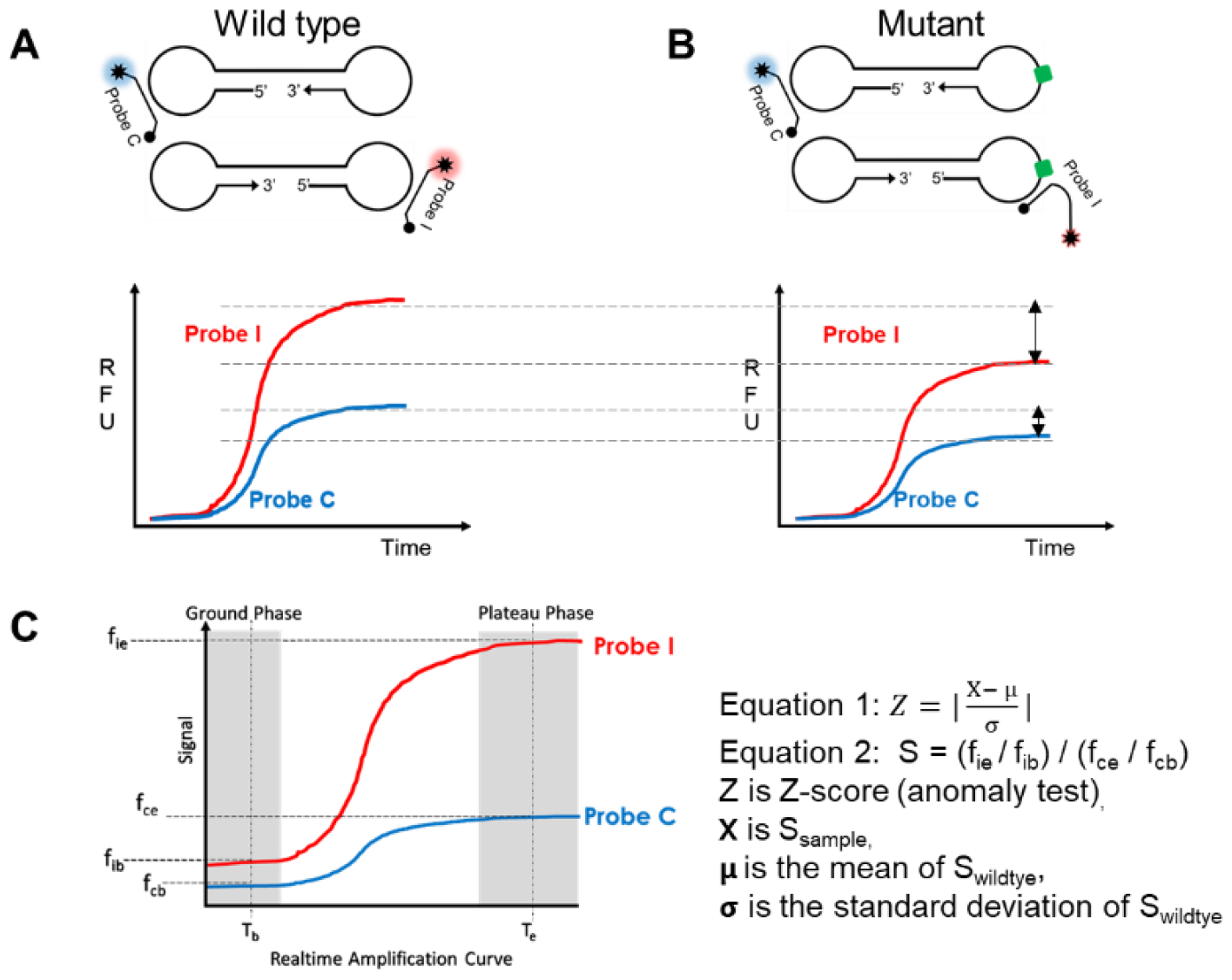
Schematic diagram of dual probe based mutation detection. (A-B) Hybridization of probes to the amplicon or intermediary amplification product obtained using isothermal amplification method like LAMP. The signals from the probes can deviate depending on the presence or absence of mutation or template concentration. Mutation is determined by deviation of calibrated signals from sample compared to that from wild type. The raw data signal is related to the amount of amplified product and hybridization efficiency between target and probe. Lower panel: an example by real time detection. WT: Wild type; Probe I: “Indicator” probe; Probe C: “Calibrator” probe. (C) A plot of dual probe detection using real-time readout in isothermal amplification. X-axis is time; Y-axis is signal; T_b_: baseline or background signal time point; T_e_: plateau phase signal end time point. Right panel: Equations for signal calibration. Z-score is used to determine presence or absence of mutation.

## MATERIALS AND METHODS

### Materials

Wild type genomic DNA of *Mycobacterium tuberculosis* strain H37Ra was obtained from ATCC (Cat#25177DQ) and quantified by Nanodrop spectrophotometer (Thermo Fisher Scientific). DNA fragments with wild type or mutation MTB DNA sequences were synthesized and cloned into pUC19 plasmid by GenScript (Figure 2). Concentrations of these plasmids are quantified by Bio-Rad ddPCR platform following the user guide. NATtrol MTB verification panel was purchased from ZeptoMetrix (Cat# NATMTBP-C).

**Figure 2:**
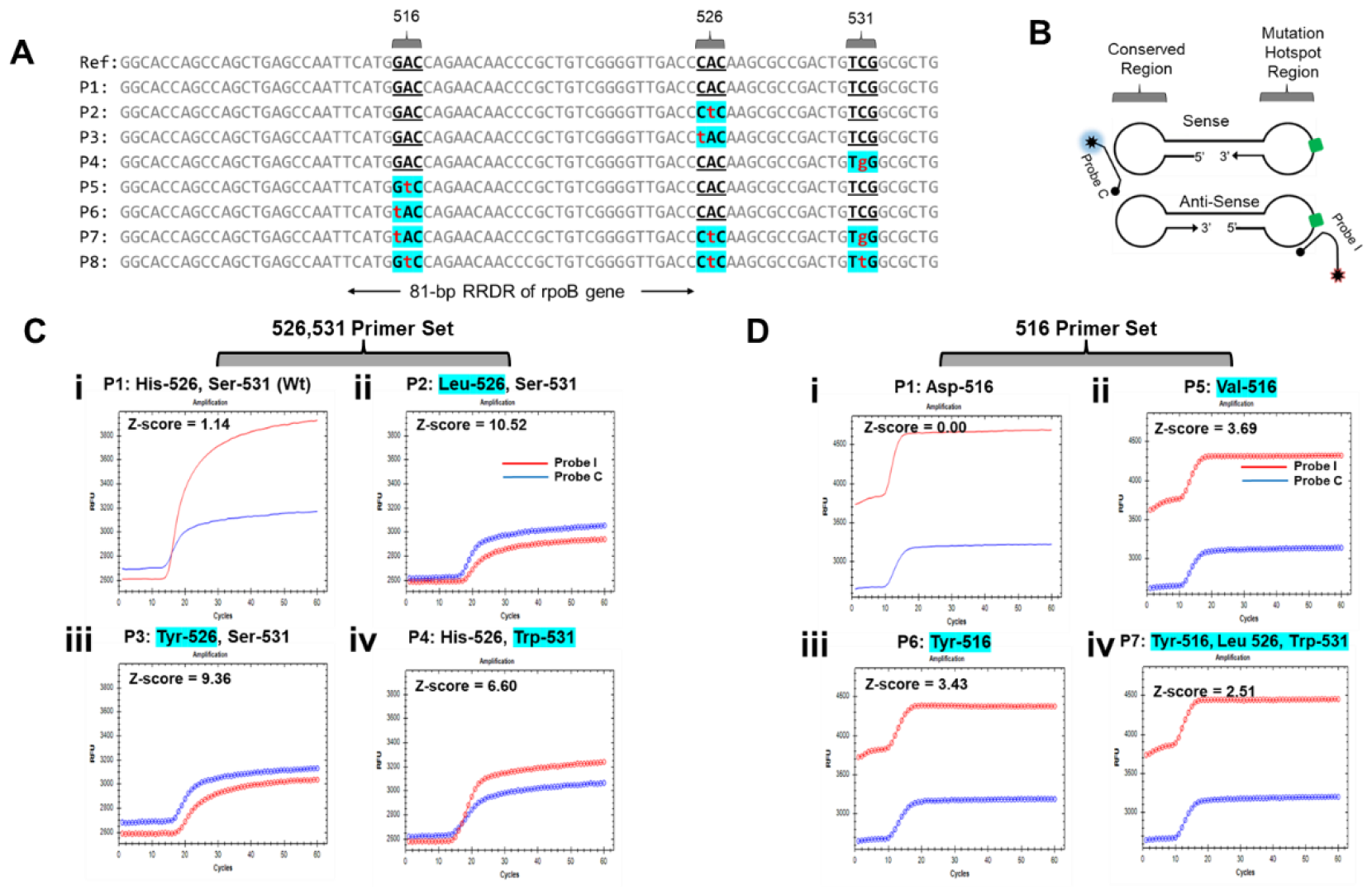
RIF-drug resistance related mutation sites and their detection by the dual probe method. (A) Sequences of plasmid inserts used for the study. The mutation sites considered in the study are highlighted in blue and the mutations are denoted in small alphabet in red. (B) Probe C and Probe I binding region RIF drug resistance detection using dual probe based strategy in LAMP. (C) Mutation detection at 526 and 531 sites using dual probe method. Fluorescence signals from the calibrator (Probe C) and indicator (Probe I) probes in presence of plasmid samples: (i) Wild type plasmid (His-526, Ser-531). (ii) P2 plasmid (Leu-526, Ser-531). (iii) P3 plasmid (Tyr-526, Ser-531). (iv) P4 plasmid (His-526, Trp-531). (D) Mutation detection at site 516 using dual probe method. Fluorescence signals from the calibrator (Probe C) and indicator (Probe I) probes in presence of plasmid samples: (i) Wild type plasmid (Asp-516). (ii) P5 plasmid (Val-516). (iii) P6 plasmid (Tyr-516). (iv) P7 plasmid (Tyr-516, Leu-526, Trp-531). X-axis is time (min), Y-axis is fluorescence signal (RFU). Z-score is labeled in each sample’s dual probe detection plot.

### Isothermal amplification and signal collection

4X-Lyo-ready LAMP mix from Meridian BioSciences (cat#MDX097) was used for the LAMP reactions. HPLC-purified primers and probes were purchased from either IDT or Sigma Aldrich, and resuspended using TE buffer (Sigma Aldrich, cat# 93283) (Supplementary Table S1). LAMP reaction was performed in a final 15 μL reaction mixture consisting of 1X-Lyo-ready LAMP mix, and 1.6 μM (each) of the primers FIP and BIP, 0.2 μM (each) of the primers F3 and B3, 0.8 μM (each) of primers LF and LB, 0.4 μM (each) of the calibrator probe (Probe C) and the indicator probe (Probe I), 8.0 mM MgSO_4_ and target nucleic acid template either genomic DNA or plasmids. Target amplification and real-time signal collection were carried out at 66°C for 60 mins on CFX96 Touch Real-Time PCR Detection System from Bio-Rad.

### Zeptometrix MTB verification panel

Samples from the Zeptometrix MTB verification panel were extracted prior to LAMP assay as described in detail below: 200μL of the wild type strain and 600μL of the RIF-resistance strain were centrifuged at 12000 rpm for 5 minutes. The pellet obtained after discarding the supernatant was resuspended in 200μL TE buffer, and then heated at 95°C for 30minutes and cooled to room temperature. Genomic DNA from the heat lysed ZeptoMetrix MTB verification panel samples was purified and eluted in 40μL TE using Oligo Clean & Concentrator (Cat# D4061, Zymo Research, CA, USA). 5μL of extracted DNA was used as input for the LAMP assay.

### Statistical Analysis

Realtime fluorescence signals obtained were divided into different regions as shown in Figure1C. Z-scores calculated based on Equation 1 and Equation 2 were used to determine the sample type as wild type or mutant.

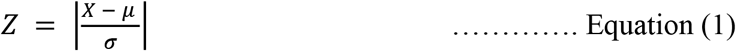

where,

X: S_sample_

μ: S_wt_

σ: standard deviation of S_wt_

the values of S for the wild type reference and test samples are calculated using the following equation,

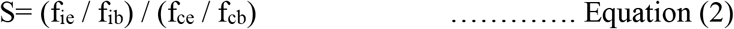

where,

f_cb_: Baseline signal of “calibrator” probe

f_ce_: End point signal of “calibrator” probe

f_ib_: Baseline signal of “indicator” probe

f_ie_: End point signal of “indicator” probe

RFU values at 10 minutes were designated as baseline signals for both the probes (f_cb_ and f_ib_) since amplification was observed after 10 minutes in most cases. As for the endpoint signals for both the probes (f_ce_ and f_ie_), RFU values at 40 minutes were selected since most signals reached plateau after 30 minutes.

A threshold value: Z_T_ for each primer set, is determined by the S values from the wild type. A Z-score greater than the threshold value (e.g., Z > Z_T_) indicates presence of mutation and a Z-score value is less than or equal to the threshold value (e.g., Z ≤ Z_T_) indicates the absence of mutation.

## RESULTS

### Dual probe method principle and statistical model for mutation detection

Isothermal techniques like LAMP that despite having rapid processivity encounter the disadvantage of being unable to provide single nucleotide differentiation due to low-stringency for prime annealing at defined temperature, typically below 68 °C.^15^ Anderson et al in their mini review have highlighted some attempts at designing quantitative LAMP assay by using different primer modification techniques.^15^ However, many methods do not address the variations in signals caused by background signals, reaction conditions, and low target copy number.^16^ To address these signal variations, we propose a dual probe method for mutation detection using isothermal amplification technology. In this method (Figure 1A-B), one probe detects a conserved region of the target nucleic acid and acts as a calibrator (Probe C), the other probe binds the region of mutation in the target nucleic acid sequence and acts as the mutation indicator (Probe I). Both Probe C and Probe I are designed such that they are complimentary to the wild type sequence. In theory, a probe complimentary to the wild type sequence, like Probe I, would give reduced or no signal in presence of the mutated sequence. However, such signal reduction or loss could be attributed to many other factors such as reaction inhibition, background noise, or a failed reaction. As Probe C binds to the same amplicon as Probe I, the signal from Probe C would be similarly affected by the reaction conditions as Probe I. This makes Probe C the ideal internal signal calibrator, to address the variations in signals due to various factors.

Since the dual probe method, takes into consideration the fluorescence signals of both the calibrator and indicator probe to determine the presence or absence of the mutation in the sample (Figure 1A-B), a statistical method model was used to process the signals. To choose a proper statistical model, we first sought to consider all the possible signal relationships between Probe I and Probe C for mutation detection (See details in Figure S1-2). To ensure the mutation detection is not affected by phenomena such as reaction inhibition, background noise, and guided by the shape of signal curves between Probe I and Probe C (Figure 1A 1B & Figure S1), our statistical model considers both the baseline signals in the ground phase and the end point signals in the plateau phase of the amplification curve. Probe I and Probe C end-point signal are normalized by their own ground phase and endpoint signals (f_ie_ / f_ib_ and f_ce_ / f_cb_ in Figure 1C) before we calculate the value of S (the ratio of Probe I to Probe C, Equation 2 in Methods). Z-score which measures how many standard deviations S_sample_ (test sample) is away from the mean of S_wt_ (wildtype samples) was finally calculated. A threshold or cut-off value of Z_T_ for Z-score was determined using reference wildtype cohort samples, any sample with Z-score greater than Z_T_ will be considered as outliers or be called mutants (see definitions and equations in Methods).

### Dual probe method doesn’t compromise the sensitivity of the assay

To test the feasibility of our dual probe mutation detection method, we designed LAMP primers to amplify RRDR region of *rpoB*, and two molecular beacon probes with Probe C binding a conserved region and Probe I binding the mutation hotspot region in the amplicon. Generally, in LAMP systems, the probes and loop primer pair of LAMP are designed in the same single-stranded region in dumbbell amplicon structure (Figure 1A), which could exhibit a competitive hybridization relationship in reaction. Introducing a second probe into LAMP reaction could potentially interfere with the efficiency of the conventional one probe-based LAMP assay. To evaluate this, different concentrations of the reference ATCC genomic DNA were tested against each probe individually and together. As seen in Figure S3, the target detection efficiency of the assay in presence of either only calibrator probe or only indicator probe or in presence of both the calibrator and the indicator probe is similar with assay sensitivity at ∼100 copies/reaction.

Another potential drawback in using probe-based detection is that the fluorescence signal can be affected by the template concentration. To evaluate if the internal calibration Probe C used in our dual probe method can help overcome this drawback, we investigated signal amplification curves with wild type and mutant sequences (Figure S4) at concentrations 37-4000 copies/reaction. As expected, fluorescence signal variations occurred across different reactions and sample concentrations. But more excitingly, both the calibrator and indicator probes signals were affected proportionally (Figure S4). In summary, using the calibrator probe signal as ruler for the indicator probe makes the system robust and independent of template concentration and other system variables.

Also, many probe-based detection methods designed to detect specific mutation sometimes cannot distinguish between wild type and other substitution mutations at the same site since the type of substitution can affect the probe binding efficiency and in turn the amplification signal. For instance, RIF drug resistance is linked to both Ser531Leu and Ser531Trp mutations. Probes designed to detect the more commonly occurring Ser531Leu might fail to distinguish between wild type and Ser531Trp. To evaluate if the dual probe method can be used to detect different mutations irrespective of the type of substitutions, we investigated contrived samples with different mutations Ser531Leu and Ser531Trp. The results in Figure S4 show that dual probe method could distinguish both Ser531Leu and Ser531Trp mutations from wild type.

### Dual probe method performance in contrived samples

To systematically evaluate the MTB RIF drug resistance detection capability of the dual probe method, we obtained eight plasmids with inserts containing major mutations at the 516, 526 and 531 codons of the RRDR regions of *rpoB* (Figure 2A). Two primer sets for LAMP were designed: one primer set for 526, 531 mutation detection, the other set for 516 mutation detection. A reference wild type training data set (N=12 for both 526, 531 primer set, and 516 primer set) to determine Z-score threshold (Z_T_) was obtained using MTB ATCC DNA (Table S2 and S3). Plasmids carrying the wild type sequence and different types and combinations of mutations were then tested with either the 526, 531 primer set (N= 47) or the 516 primer set (N=80). The fluorescence values at baseline and endpoint for both the calibrator and indicator probes used to calculate the Z-score values for the samples are given in Tables S4-S5.

Figure 2B shows the binding sites for Probe C and Probe I on the sense and anti-sense strands of the amplicon. Figure 2C and 2D show example amplification curves of some of the samples tested (samples in bold in Table S4 and S5). In the 526, 531 primer set shown in Figure 2C, consistent fluorescence signals from Probe C were observed irrespective of presence or absence of mutations. This consistency in Probe C signal (blue amplification curves) is as expected since it is designed to bind a conserved sequence on the amplicon and remains unaffected by the wild type or mutant sequences. The fluorescence sequence from Probe I on the other hand is fully complementary to the wildtype sequence (Figure 2B). The changes in Probe I signal intensities (red amplification curves) observed between wild type and mutation sequence can be attributed to the differential binding efficiency of Probe I to the target region (Figure 2B-C). The results also show that mutations located in close proximity (example codon 526 and 531) could be detected using same set of primers and probes. Similar trends for Probe C and Probe I signals could be observed in 516 primer set as seen Figure 2D.

The 47 samples tested with the 526, 531 primer set were comprised of different concentrations of wild type sequences and RIF mutations at codons 526 and 531. Among the 47 samples tested, 7 samples were considered invalid due to failed reactions with template concentrations falling below the reaction sensitivity, as indicated in Figure S2. Using the Z_T_ of 5.5 obtained from training data set, the remaining 40 samples were differentiated into wild type and mutant (Figure 3A). 11 out of 11 were correctly identified as wild type and all the 29 samples with mutated sequences were identified with 100% accuracy irrespective of the concentration of the plasmid in the sample (Figure 3A 2X2 table).

**Figure 3:**
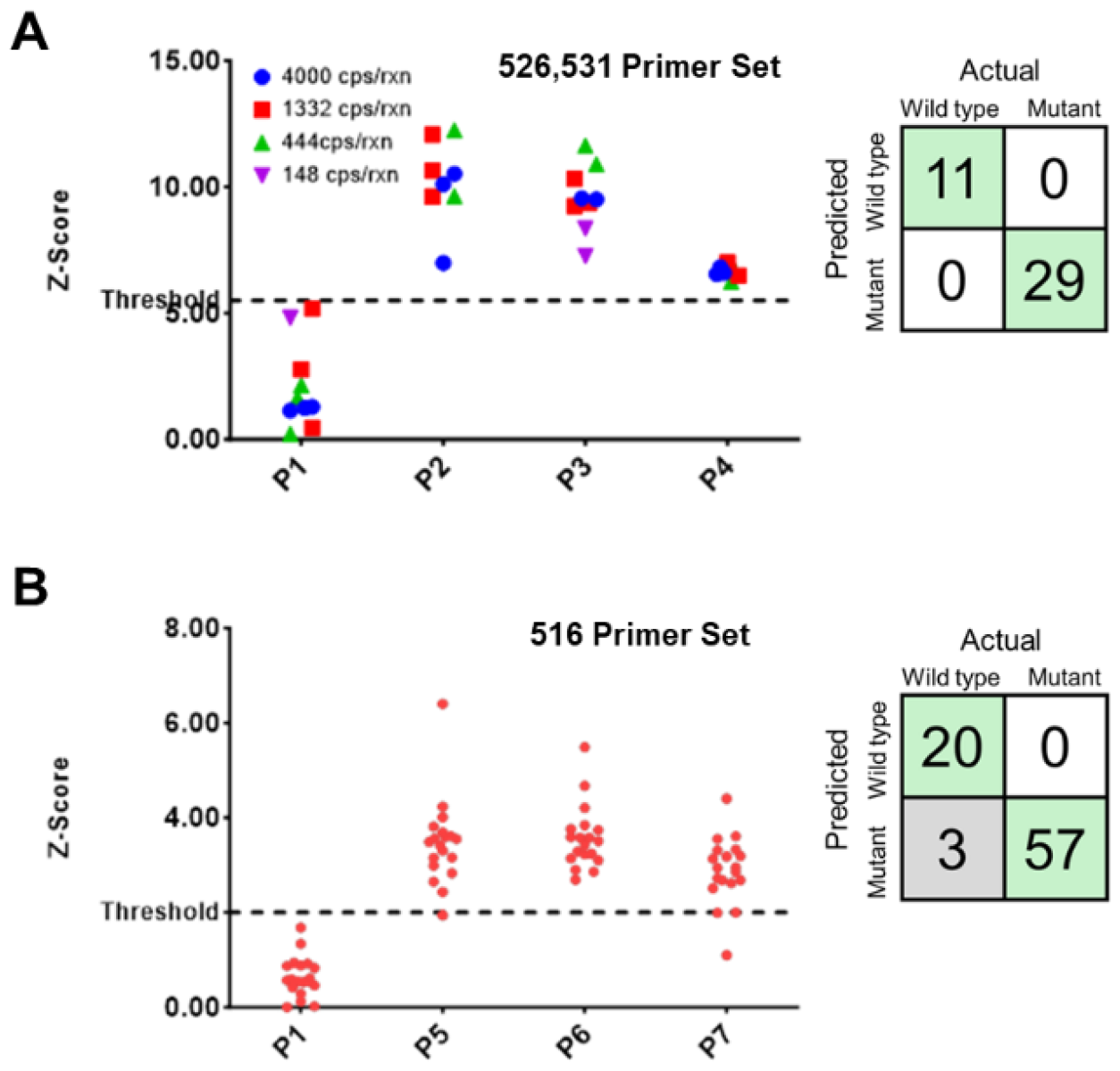
Dual probe based mutation detection performance on contrived samples. (A) Scatter plot of Z-score of 40 valid contrived samples of different genotypes at different concentrations for detection of mutations at sites 526 and 531 using the 526, 531 primer set. Initial sample concentration in reaction is labeled using different shapes. Dashed line Z-score= 5.5 is a threshold. The 2X2 table showing 100% accuracy in determining the sample type. (B) Scatter plot of Z-score of 80 valid contrived samples of different genotypes at different concentrations for detection of mutations at site 516 using the 516 primer set. Dashed line Z-score= 2 is a threshold. The 2X2 table shows 100% accuracy for wild type detection and 95% accuracy for mutation detection.

Since the template concentration did not influence the sample identification, samples at a constant concentration of 1000 copies/reaction were tested for the 516 primer set. A Z_T_ value of 2 was obtained from the training dataset using the 516 primer set. 80 samples comprising of 20 wild type sequences and 60 sequences with RIF mutations at codon 516 were tested using the assay (Figure 3B). With a threshold of 2, 20 out of 20 were correctly identified as wild type and 57 out of 60 samples with mutated sequences were identified with 95% accuracy (Figure 3B 2X2 table).

### Dual probe method validation in Zeptometrix MTB verification Panel

To further evaluate the dual probe method for MTB drug resistance detection, we obtained MTB verification panel from Zeptometrix. These samples are formulated with purified, intact bacterial cells. After extraction and purification, wildtype and mutant strains’ DNAs were tested by our 526,531 LAMP primer set and 516 LAMP primer set. Plots for reference and wild type Zeptometrix sample show Z-score below the threshold values for individual oligo sets (Z_T526,531_=5.5, Z_T 516_=2), Figure 4A i, ii and Figure 4B i and ii. The plot in Figure 4A iii, shows a Z-score value of 8.76 which is higher than the threshold value of 5.5, indicating presence of mutation at 526 and/or 531. However, the plot in Figure 4B iii, has a Z-score of 1.08 which is below the threshold value, Z_T516_=2, indicating absence of mutation at codon 516. This agrees with the Ser531Leu mutation present in the Zeptometrix RIF-resistant strain.

**Figure 4:**
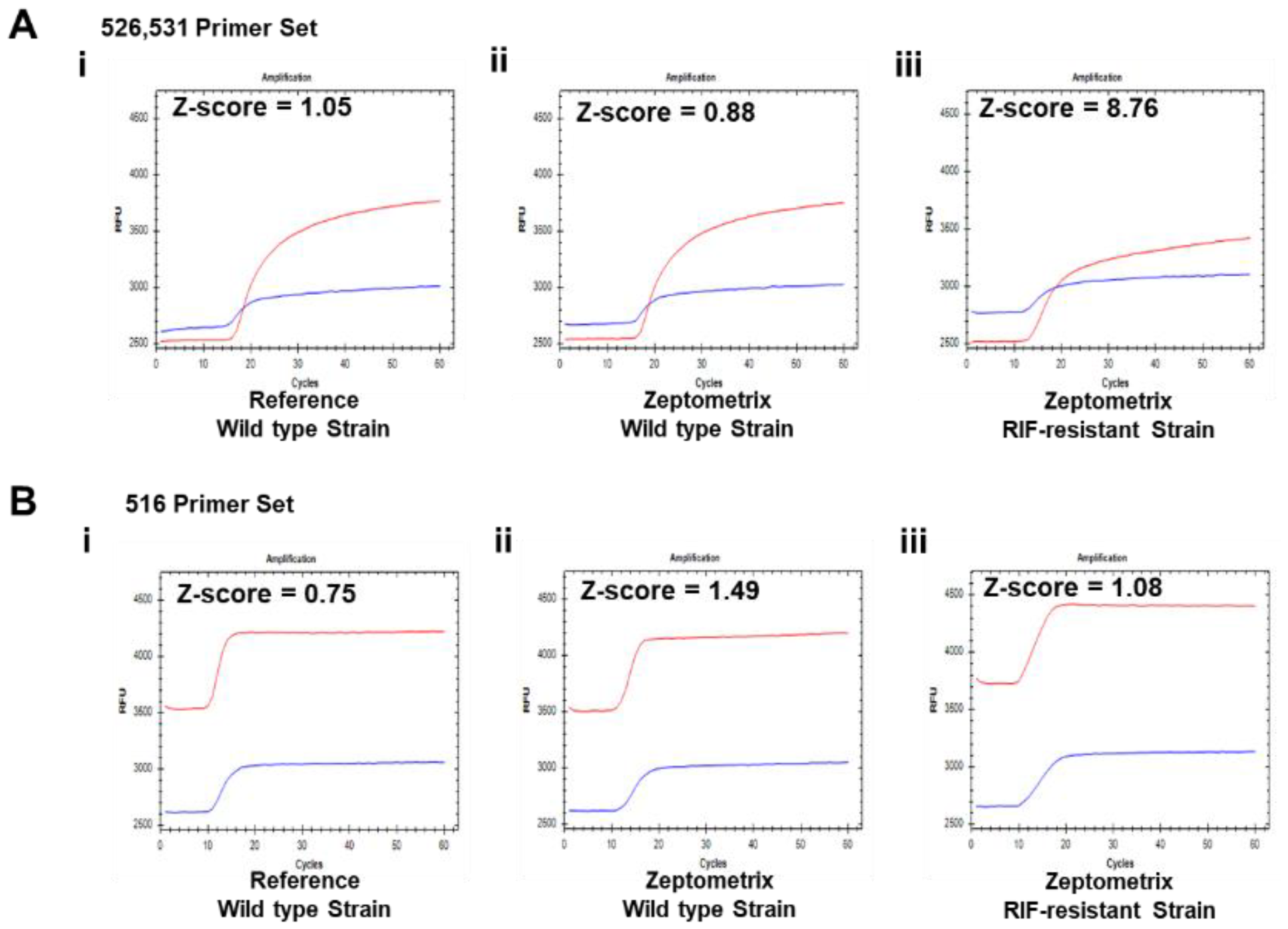
Mutation detection in MTB verification panel. (A) Fluorescence signals from the calibrator (Probe C) and indicator (Probe I) probes (designed to detect mutations at sites 526 and 531) in presence of MTB verification panel samples: (i)Reference Wild type plasmid (His-526, Ser-531). (ii) Zeptometrix Wild type strain (iii) Zeptometrix Mutant type strain carrying mutation Ser531Leu. (B) Fluorescence signals from the calibrator (Probe C) and indicator (Probe I) probes (designed to detect mutations at site 516) in presence of MTB verification panel samples: (iv)Reference Wild type plasmid (His-526, Ser-531). (v) Zeptometrix Wild type strain (vi) Zeptometrix Mutant type strain carrying mutation Ser531Leu. X-axis is time (min), Y-axis is fluorescence signal (RFU). Z-score is labeled in each sample’s dual probe detection plot.

## DISCUSSIONS

Many assays designed for drug resistance related mutation detection can be relatively expensive, require sophisticated instruments, impacted by inhibitors in sample resulting in false-negative results. This makes using such methods less than ideal for high threat regions. The dual probe method proposed here has simple ingredients, is affordable, rapid, doesn’t require sophisticated instruments. The incorporation of a probe as an internal control serves to mitigate signal variations from template concentration or system variables (e.g., instrument fluorescence readings) or reaction conditions (e.g., inhibitions from sample matrix). This, in turn, contributes to a reduction in both false positive and false negative rates. Many systems require the use of multiple probes in order to detect each type of mutation, which is necessary when studying different substitutions at given mutation site. However, in cases where mutation irrespective of the type of the substitution or an emerging new mutation leading to drug resistance are encountered, the current dual probe method is a simpler design that allows for rapid mutation detection.

Many point-of-care colorimetric based LAMP reactions have already been reported. Our method, however, overcomes one of the major drawbacks related to specificity encountered in these LAMP reactions. First, using molecular beacon ensures target specificity. Second, the incorporation of the calibrator probe ensures that the mutation detection is not affected by template concentration and other system variables. The probes in our method detect target regions on the same amplicon allowing us to use single primer set for amplification. Using this assay design, we were able to identify mutations at codon 516, 526 and 531, that cover more than 90% RIF related drug resistance cases reported, with 100% accuracy for mutations at 526 and 531 sites and 95% accuracy for mutations at site 516 in the evaluated sample set. We were also able to distinguish between the wild type and mutation strain from the Zeptometrix verification samples that are used as controls for assay development.

In conclusion, the dual probe technology holds great promise for mutation detection. Specifically, when used with isothermal nucleic acid amplification, its rapidity, affordability, and suitability for resource-limited settings make it an attractive alternative to conventional PCR methods. By enabling timely detection of drug-resistant strains, this technology has the potential to guide appropriate treatment decisions, limit the spread of resistance, and improve patient outcomes in infectious disease control programs. It is also essential to note that the dual probe concept has its limitations. It may not be able to differentiate drug-resistance conferring mutations from silent mutations at the same locations ^17,18^, and it may not be able determine whether the samples are a heterozygous pool or homozygous pool.

## DATA AVAILABILITY

All data generated or analyzed during this study are included in this published article and its Supplementary Information files.

## ACKNOWLEDGEMENTS

We thank Dr. Wolfgang Pieken, Dr. Yali Sun, and Dr. Macy Veling for critical discussion of this work.

## AUTHOR CONTRIBUTIONS

Z.L. conceived the study. Z.L. and Y.T. supervised the project. N.N., M.M. and Z.L. performed the key experiments. N.N. and Z.L. carried out the analysis and drafted the paper. All authors reviewed the paper.

