## Supplemental Data for "A Novel Dual Probe-based Method for Mutation Detection using Isothermal Amplification"

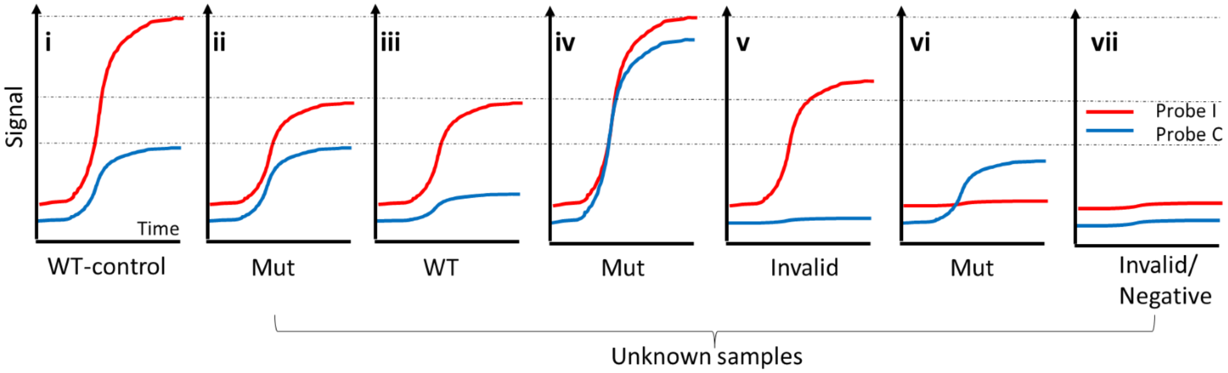


**Supplemental Figure 1**. Schematic representation of possible results using the dual-probe method (using real-time readout as an example). (i) Realtime amplification curve of Wild type control sample. (ii) A mutant sample in which calibrator probe signal does not change while indicator probe signal dramatically drops due to effects of mutations on the indicator probe-target hybridization efficiency. (iii) A wild type in which signals from calibrator probe and indicator probe synchronously decrease. This pattern might result from relatively low concentration of the sample, or high non-specific product, low reaction activity. The sample type identification is not affected since, calibrated score does not change. (iv) A mutant in which calibrator probe signal increases while indicator probe signal does not change. Calibrated score changes. (v) Invalid as there is no signal from endogenous reference probe (i.e., calibrator probe). The indicator probe signal could be a result of some cross contamination or non-specific amplification. (vi) A mutant that calibrator probe shows presence of target, no signal from mutation indicator probe. (vii) Negative or Invalid as neither of probes has signals. This result might be due to expired reagents, sample interference, or low concentration or absence of target nucleic acid.

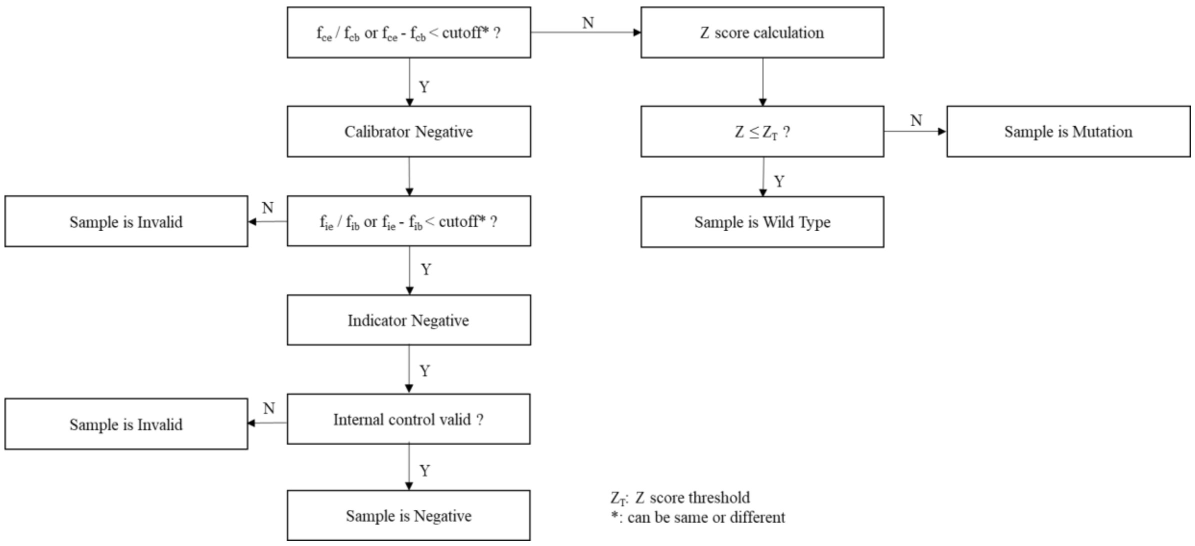


**Supplemental Figure 2**. Decision Flow chart for sample type determination

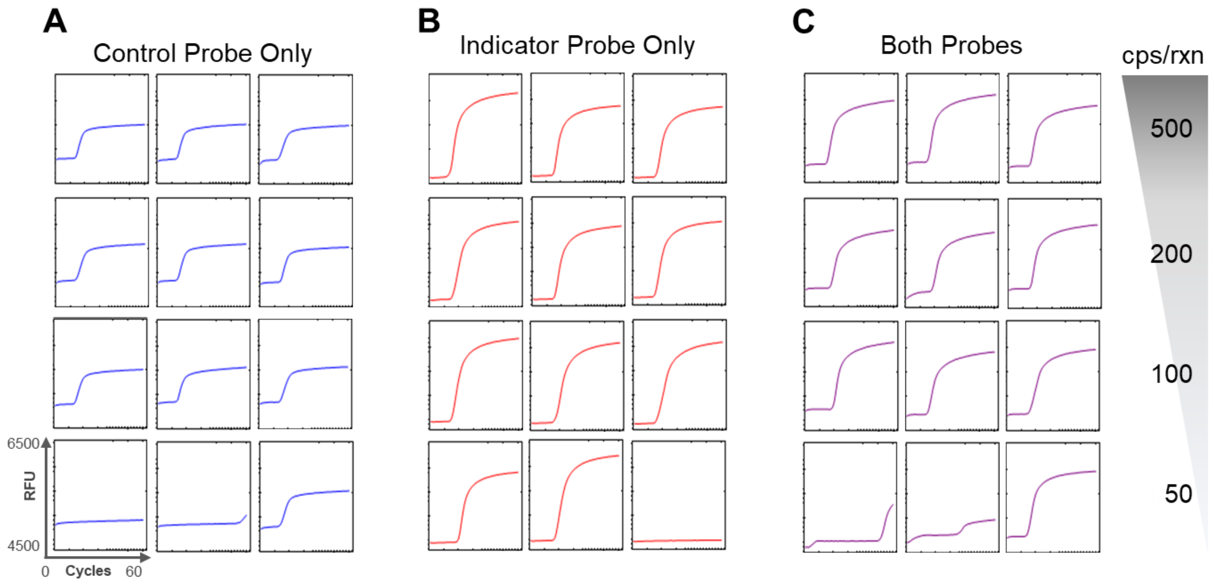


**Supplemental Figure 3. Effect of presence of two probes on efficiency of assay.** (A) Fluorescence signals from the calibrator (Probe C) probe (designed to detect mutations at sites 526 and 531) in presence of MTB reference wild type genomic DNA. (B) Fluorescence signals from the indicator (Probe I) probe (designed to detect mutations at sites 526 and 531) in presence of MTB reference wild type genomic DNA. The sensitivity of the assay lies between 100 and 50 copies/reaction irrespective of the presence of single or both probes. X-axis is time (min), Y-axis is fluorescence signal (RFU).

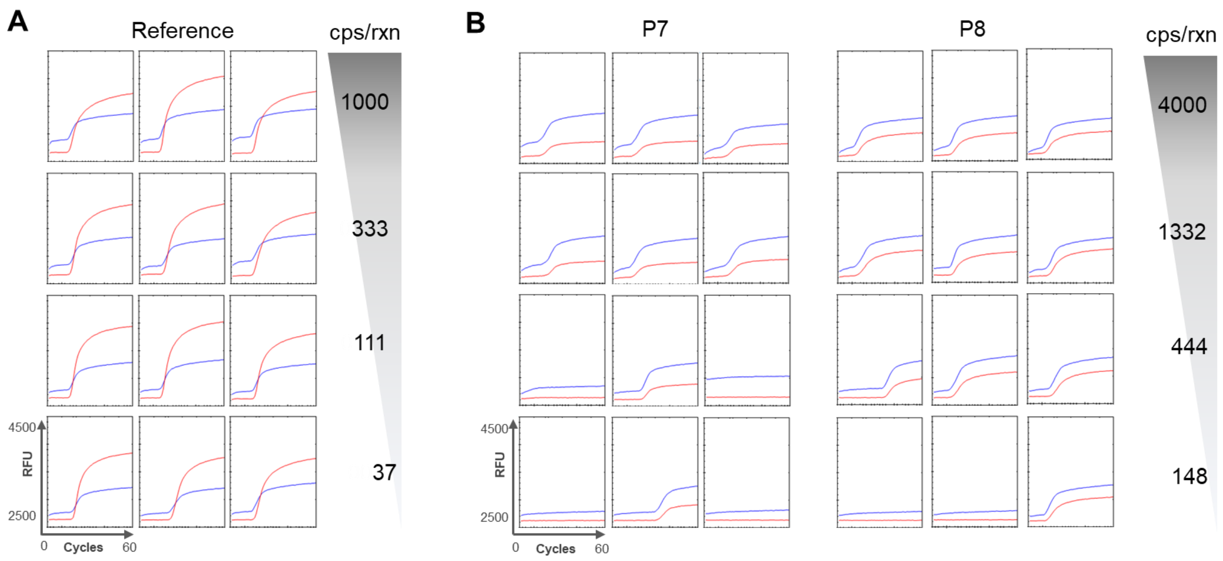


**Supplemental Figure 4. Effect of template concentration on assay efficiency.** (A) Fluorescence signals from the calibrator (Probe C) probe (designed to detect mutations at sites 526 and 531) in presence of 37-1000 copies/reaction of MTB reference wild type genomic DNA. (B) Fluorescence signals from the calibrator (Probe C) and indicator (Probe I) probes (designed to detect mutations at sites 526 and 531) in presence of contrived plasmid samples. The probe signals are similar irrespective of the template concentration. X-axis is time (min), Y-axis is fluorescence signal (RFU).

**Supplemental Table 1**. Sequences of primers and probes

| Oligo Name | Sequences (5' – 3') |
|---|---|
| $F3_{526,531}$ | AGCGGATGACCACCCAG |
| $B3_{526,531}$ | TGCACGTCGCGGACCT |
| $FIP_{526,531}$ | CTTGATCGCGGCGACCACCGGACGTGGAGGCGATCACA |
| $BIP_{526,531}$ | CAGAACAACCCGCTGTCGGCACGCTCACGTGACAGACC |
| $LF_{526,531}$ | GCCGGATGTTGATCAACG |
| $LB_{526,531}$ | CCACAAGCGCCGACTG |
| Probe $C_{526,531}$ | FAM-CGCGAGCCGGATGTTGATCAACGTCTGCTCGCG-BHQ1 |
| Probe $I_{526,531}$ | Cy5-CGCGAGACC[+C][+A][+C]AAGCGCCGACTG[+T][+C][+G]GCGCTCGCG-BHQ2* |
| $F3_{516}$ | AGCGGATGACCACCCAG |
| $B3_{516}$ | CGCTCACGTGACAGACCG |
| $FIP_{516}$ | CTTGATCGCGGCGACCACCGGACGTGGAGGCGATCACA |
| $BIP_{516}$ | GTTCTTCGGCACCAGCCAGCTGCCGACAGTCGGCGCTT |
| $LF_{516}$ | GATGTTGATCAACGTCTGCG |
| $LB_{516}$ | GAGCCAATTCATGGACCAGAA |
| Probe $C_{516}$ | Cy5-CGCGAGCCGGATGTTGATCAACGTCGCG-BHQ2 |
| Probe $I_{516}$ | FAM-CGCGACCAATTCATG[+G][+A][+C]CAGAACAACTCGCG-BHQ1* |

*: [+] is LNA modified base.

**Supplemental Table 2**. $S_{wt,526,531}$ calculation using *Mycobacterium tuberculosis* genomic DNA.

| Standards | $f_{ib}$ | $f_{ie}$ | $f_{cb}$ | $f_{ce}$ | *$S_{wt,526,531}$= ($f_{ie}$ / $f_{ib}$) / ($f_{ce}$ / $f_{cb}$) |
|---|---|---|---|---|---|
| MTB-gDNA | 2664 | 3621 | 2891 | 3323 | 1.18 |
| MTB-gDNA | 2668 | 3927 | 2903 | 3390 | 1.26 |
| MTB-gDNA | 2657 | 3658 | 2940 | 3401 | 1.19 |
| MTB-gDNA | 2662 | 3826 | 2842 | 3295 | 1.24 |
| MTB-gDNA | 2675 | 3812 | 2822 | 3260 | 1.23 |
| MTB-gDNA | 2650 | 3666 | 2910 | 3354 | 1.20 |
| MTB-gDNA | 2649 | 3855 | 2785 | 3237 | 1.25 |
| MTB-gDNA | 2650 | 3915 | 2796 | 3282 | 1.26 |
| MTB-gDNA | 2632 | 3699 | 2761 | 3212 | 1.21 |
| MTB-gDNA | 2639 | 3743 | 2762 | 3170 | 1.24 |
| MTB-gDNA | 2633 | 3615 | 2746 | 3153 | 1.20 |
| MTB-gDNA | 2634 | 3627 | 2779 | 3245 | 1.18 |

*Note: Mean (μ) $S_{wt,526,531}$ =1.22, Standard Deviation (σ) $S_{wt,526,531}$ =0.03

**Supplemental Table 3**. $S_{wt,516}$ calculation using *Mycobacterium tuberculosis* genomic DNA.

| Standards | $f_{ib}$ | $f_{ie}$ | $f_{cb}$ | $f_{ce}$ | *$S_{wt,516}$= ($f_{ie}$ / $f_{ib}$) / ($f_{ce}$ / $f_{cb}$) |
|---|---|---|---|---|---|
| MTB-gDNA | 3771 | 4603 | 2679 | 3204 | 1.02 |
| MTB-gDNA | 3739 | 4430 | 2697 | 3178 | 1.01 |
| MTB-gDNA | 3707 | 4413 | 2669 | 3131 | 1.01 |
| MTB-gDNA | 3625 | 4261 | 2667 | 3111 | 1.01 |
| MTB-gDNA | 3585 | 4295 | 2652 | 3105 | 1.02 |
| MTB-gDNA | 3614 | 4274 | 2660 | 3106 | 1.01 |
| MTB-gDNA | 3807 | 4484 | 2678 | 3136 | 1.01 |
| MTB-gDNA | 3554 | 4161 | 2659 | 3084 | 1.01 |
| MTB-gDNA | 3777 | 4353 | 2701 | 3130 | 0.99 |
| MTB-gDNA | 3677 | 4292 | 2684 | 3137 | 1.00 |
| MTB-gDNA | 3858 | 4424 | 2689 | 3160 | 0.98 |
| MTB-gDNA | 3694 | 4281 | 2684 | 3113 | 1.00 |

*Note: Mean (μ) $S_{wt,516}$ =1.01, Standard Deviation (σ) $S_{wt,516}$=0.01

**Supplemental Table 4**. Z-score for the 47 samples tested using the dual-probe method. (Mutations at 526 and 531) (Data in bold was used to create the example amplification curves in Figure 2c)

| Plasmid-ID | Concentration (cp/reaction) | $f_{ib}$ | $f_{ie}$ | $f_{cb}$ | $f_{ce}$ | $S_s$= ($f_{ie}$ / $f_{ib}$)/ ($f_{ce}$ / $f_{cb}$) | Z-Score |
|---|---|---|---|---|---|---|---|
| P1 | 4000 | 2615 | 3760 | 2705 | 3101 | 1.25 | 0.49 |
| **P1** | **4000** | **2608** | **3833** | **2702** | **3127** | **1.27** | **1.14** |
| P1 | 4000 | 2591 | 3593 | 2727 | 3128 | 1.21 | 1.29 |
| P1 | 1332 | 2585 | 3597 | 2695 | 3100 | 1.21 | 1.25 |
| P1 | 1332 | 2572 | 3724 | 2646 | 3114 | 1.23 | 0.46 |
| P1 | 1332 | 2562 | 3517 | 2608 | 3057 | 1.17 | 2.77 |
| P1 | 444 | 2572 | 3348 | 2621 | 3072 | 1.11 | 5.18 |
| P1 | 444 | 2580 | 3643 | 2696 | 3080 | 1.24 | 0.22 |
| P1 | 444 | 2583 | 3596 | 2642 | 3065 | 1.20 | 1.65 |
| P1 | 148 | 2589 | 3523 | 2684 | 3075 | 1.19 | 2.14 |
| P1 | 148 | 2553 | 3288 | 2587 | 2976 | 1.12 | 4.81 |
| P2 | 4000 | 2593 | 2931 | 2612 | 2997 | 0.99 | 10.17 |
| **P2** | **4000** | **2590** | **2906** | **2621** | **3012** | **0.98** | **10.52** |
| P2 | 4000 | 2594 | 3191 | 2648 | 3059 | 1.07 | 6.99 |
| P2 | 1332 | 2580 | 2933 | 2628 | 3028 | 0.99 | 10.11 |
| P2 | 1332 | 2583 | 2958 | 2629 | 3013 | 1.00 | 9.62 |
| P2 | 1332 | 2595 | 2819 | 2660 | 3086 | 0.94 | 12.08 |
| P2 | 444 | 2586 | 2853 | 2632 | 2984 | 0.97 | 10.65 |
| P2 | 444 | 2582 | 3010 | 2637 | 3079 | 1.00 | 9.64 |
| P2 | 444 | 2588 | 2790 | 2642 | 3055 | 0.93 | 12.25 |
| P2 | 148 | 2578 | 2581 | 2637 | 2650 | - | Invalid |
| P2 | 148 | 2575 | 2575 | 2617 | 2635 | - | Invalid |
| P2 | 148 | 2581 | 2574 | 2636 | 2644 | - | Invalid |
| P3 | 4000 | 2590 | 2991 | 2608 | 3006 | 1.00 | 9.50 |
| P3 | 4000 | 2589 | 3003 | 2646 | 3066 | 1.00 | 9.51 |
| P3 | 4000 | 2649 | 3059 | 2988 | 3115 | - | Invalid |
| P3 | 1332 | 2588 | 3016 | 2642 | 3077 | 1.00 | 9.55 |
| **P3** | **1332** | **2591** | **2992** | **2691** | **3091** | **1.01** | **9.36** |
| P3 | 1332 | 2586 | 3042 | 2640 | 3080 | 1.01 | 9.24 |
| P3 | 444 | 2594 | 2922 | 2666 | 3062 | 0.98 | 10.33 |
| P3 | 444 | 2578 | 2847 | 2647 | 3084 | 0.95 | 11.64 |
| P3 | 444 | 2584 | 2904 | 2655 | 3087 | 0.97 | 10.90 |
| P3 | 148 | 2590 | 2591 | 2773 | 2788 | 0.99 | 9.77 |
| P3 | 148 | 2571 | 3060 | 2626 | 3033 | 1.03 | 8.36 |
| P3 | 148 | 2582 | 3017 | 2854 | 3150 | 1.06 | 7.26 |
| P4 | 4000 | 2601 | 3233 | 2622 | 3022 | 1.08 | 6.48 |
| P4 | 4000 | 2599 | 3238 | 2647 | 3064 | 1.08 | 6.55 |
| P4 | 4000 | 2585 | 3208 | 2636 | 3059 | 1.07 | 6.82 |
| **P4** | **1332** | **2582** | **3190** | **2629** | **3022** | **1.08** | **6.60** |
| P4 | 1332 | 2585 | 3216 | 2629 | 3072 | 1.06 | 7.01 |
| P4 | 1332 | 2587 | 3239 | 2644 | 3086 | 1.07 | 6.69 |
| P4 | 444 | 2587 | 3262 | 2644 | 3094 | 1.08 | 6.49 |
| P4 | 444 | 2577 | 3206 | 2633 | 3050 | 1.07 | 6.66 |
| P4 | 444 | 2590 | 3267 | 2663 | 3100 | 1.08 | 6.25 |
| P4 | 148 | 2578 | 2577 | 2634 | 2651 | - | Invalid |
| P4 | 148 | 2576 | 2577 | 2633 | 2654 | - | Invalid |
| P4 | 148 | 2573 | 2568 | 2645 | 2659 | - | Invalid |

**Supplemental Table 5**. Z-score for the 80 samples tested using the dual-probe method. (Mutations at 516) (Data in bold was used to create the example amplification curves in Figure 2d)

| Plasmid-ID | $f_{ib}$ | $f_{ie}$ | $f_{cb}$ | $f_{ce}$ | $S_s = (f_{ie} / f_{ib}) / (f_{ce} / f_{cb})$ | Z-Score |
|---|---|---|---|---|---|---|
| P1 | 3888 | 4734 | 2664 | 3179 | 1.02 | 1.08 |
| P1 | 3790 | 4510 | 2648 | 3126 | 1.01 | 0.12 |
| P1 | 3847 | 4613 | 2661 | 3149 | 1.01 | 0.54 |
| P1 | 3867 | 4670 | 2656 | 3162 | 1.01 | 0.61 |
| P1 | 3877 | 4688 | 2668 | 3170 | 1.02 | 0.87 |
| P1 | 3803 | 4539 | 2653 | 3125 | 1.01 | 0.53 |
| P1 | 3798 | 4559 | 2661 | 3138 | 1.02 | 0.92 |
| P1 | 3811 | 4531 | 2661 | 3123 | 1.01 | 0.54 |
| P1 | 3683 | 4383 | 2647 | 3093 | 1.02 | 0.94 |
| P1 | 3585 | 4237 | 2626 | 3049 | 1.02 | 0.88 |
| P1 | 3872 | 4680 | 2663 | 3174 | 1.01 | 0.57 |
| P1 | 3586 | 4323 | 2632 | 3086 | 1.03 | 1.68 |
| P1 | 3861 | 4609 | 2666 | 3164 | 1.01 | 0.02 |
| P1 | 3903 | 4655 | 2672 | 3155 | 1.01 | 0.28 |
| **P1** | **3875** | **4669** | **2687** | **3217** | **1.01** | **0.00** |
| P1 | 3862 | 4636 | 2682 | 3181 | 1.01 | 0.46 |
| P1 | 3839 | 4638 | 2675 | 3188 | 1.01 | 0.59 |
| P1 | 3852 | 4614 | 2669 | 3161 | 1.01 | 0.42 |
| P1 | 3922 | 4742 | 2679 | 3184 | 1.02 | 0.83 |
| P1 | 3732 | 4477 | 2658 | 3115 | 1.02 | 1.34 |
| P5 | 3777 | 4309 | 2672 | 3148 | 0.97 | 2.90 |
| P5 | 3715 | 4225 | 2659 | 3121 | 0.97 | 2.83 |
| P5 | 3737 | 4216 | 2658 | 3127 | 0.96 | 3.61 |
| P5 | 4110 | 4492 | 2699 | 3198 | 0.92 | 6.40 |
| P5 | 3823 | 4356 | 2670 | 3171 | 0.96 | 3.55 |
| P5 | 3960 | 4543 | 2688 | 3234 | 0.95 | 4.01 |
| **P5** | **3892** | **4439** | **2682** | **3194** | **0.96** | **3.69** |
| P5 | 3949 | 4547 | 2699 | 3189 | 0.97 | 2.43 |
| P5 | 3912 | 4453 | 2675 | 3184 | 0.96 | 3.81 |
| P5 | 3692 | 4188 | 2648 | 3114 | 0.96 | 3.16 |
| P5 | 3712 | 4203 | 2658 | 3125 | 0.96 | 3.30 |
| P5 | 3751 | 4275 | 2663 | 3157 | 0.96 | 3.42 |
| P5 | 3705 | 4241 | 2647 | 3118 | 0.97 | 2.65 |
| P5 | 3829 | 4374 | 2688 | 3229 | 0.95 | 4.23 |
| P5 | 3809 | 4360 | 2680 | 3197 | 0.96 | 3.56 |
| P5 | 3869 | 4545 | 2676 | 3205 | 0.98 | 1.94 |
| P5 | 3760 | 4297 | 2671 | 3157 | 0.97 | 2.99 |
| P5 | 3829 | 4359 | 2681 | 3178 | 0.96 | 3.49 |
| P5 | 3758 | 4286 | 2654 | 3156 | 0.96 | 3.59 |
| P5 | 3700 | 4231 | 2652 | 3142 | 0.96 | 3.15 |
| P6 | 3891 | 4335 | 2713 | 3187 | 0.95 | 4.41 |
| P6 | 3807 | 4343 | 2683 | 3197 | 0.96 | 3.74 |
| P6 | 3778 | 4268 | 2671 | 3146 | 0.96 | 3.59 |
| **P6** | **3851** | **4378** | **2692** | **3185** | **0.96** | **3.43** |
| P6 | 3880 | 4430 | 2694 | 3207 | 0.96 | 3.58 |
| P6 | 3892 | 4422 | 2666 | 3166 | 0.96 | 3.76 |
| P6 | 3791 | 4323 | 2669 | 3160 | 0.96 | 3.29 |
| P6 | 3988 | 4498 | 2678 | 3175 | 0.95 | 4.21 |
| P6 | 3793 | 4285 | 2670 | 3144 | 0.96 | 3.58 |
| P6 | 3982 | 4369 | 2673 | 3139 | 0.93 | 5.49 |
| P6 | 3643 | 4140 | 2661 | 3122 | 0.97 | 2.86 |
| P6 | 3709 | 4194 | 2671 | 3134 | 0.96 | 3.23 |

| | | | | | | |
|---|---|---|---|---|---|---|
| P6 | 3672 | 4172 | 2656 | 3126 | 0.97 | 3.10 |
| P6 | 3823 | 4305 | 2717 | 3187 | 0.96 | 3.51 |
| P6 | 3870 | 4394 | 2688 | 3193 | 0.96 | 3.84 |
| P6 | 3808 | 4337 | 2672 | 3153 | 0.97 | 3.14 |
| P6 | 4015 | 4535 | 2714 | 3245 | 0.94 | 4.67 |
| P6 | 3762 | 4309 | 2656 | 3133 | 0.97 | 2.69 |
| P6 | 3752 | 4308 | 2665 | 3160 | 0.97 | 2.90 |
| P6 | 3666 | 4168 | 2643 | 3118 | 0.96 | 3.24 |
| P6 | 3706 | 4223 | 2665 | 3121 | 0.97 | 2.55 |
| P7 | 3667 | 4179 | 2676 | 3112 | 0.98 | 2.00 |
| P7 | 3634 | 4128 | 2664 | 3138 | 0.96 | 3.19 |
| P7 | 3823 | 4345 | 2682 | 3179 | 0.96 | 3.61 |
| P7 | 3660 | 4284 | 2657 | 3174 | 0.98 | 1.99 |
| P7 | 3759 | 4304 | 2673 | 3152 | 0.97 | 2.68 |
| **P7** | **3769** | **4318** | **2657** | **3128** | **0.97** | **2.51** |
| P7 | 3712 | 4242 | 2656 | 3125 | 0.97 | 2.68 |
| P7 | 3806 | 4344 | 2665 | 3144 | 0.97 | 2.95 |
| P7 | 3747 | 4286 | 2653 | 3123 | 0.97 | 2.62 |
| P7 | 3692 | 4152 | 2667 | 3125 | 0.96 | 3.55 |
| P7 | 3597 | 4066 | 2653 | 3107 | 0.97 | 3.13 |
| P7 | 3588 | 4047 | 2665 | 3102 | 0.97 | 2.85 |
| P7 | 3725 | 4172 | 2682 | 3167 | 0.95 | 4.40 |
| P7 | 3684 | 4175 | 2668 | 3141 | 0.96 | 3.31 |
| P7 | 3678 | 4181 | 2670 | 3147 | 0.96 | 3.18 |
| P7 | 3648 | 4114 | 2645 | 3099 | 0.96 | 3.33 |
| P7 | 3688 | 4193 | 2662 | 3127 | 0.97 | 2.94 |
| P7 | 3801 | 4351 | 2658 | 3135 | 0.97 | 2.72 |
| P7 | 3775 | 4440 | 2645 | 3137 | 0.99 | 1.10 |
